# Pro-inflammatory response of non-inflammatory Classical monocytes stimulated with LPS *in vitro*

**DOI:** 10.1101/2020.05.04.077537

**Authors:** Claudio Karsulovic, Fabian Tempio, Mercedes Lopez, Julia Guerrero, Annelise Goecke

## Abstract

**BACKGROUND:** CD14 (Monocyte identifying Toll-Like Receptor) and CD16 (FcyRIII co-receptor, marker of inflammatory monocytes) were used to define 3 subpopulations of circulating monocytes with different attributes in terms of inflammatory and phagocytic capabilities. There are contradictory reports regarding response of circulating monocytes to pro-inflammatory or non-inflammatory stimuli *in vitro*. Here we aimed to analyze the phenotypic changes in circulating monocytes when stimulated with pro and non-inflammatory stimuli.

**METHODS:** Whole blood from 9 healthy donors was extracted and studied. Monocyte subpopulations were directly measured using flow cytometry with PBMC Ficoll extraction method. Pro-inflammatory interleukin IL-1β was measured by intracellular cytometry. Whole blood-extracted monocytes were stimulated using LPS and IL-4 as previously described. Changes against non-stimulated (N-S) populations were statistically analyzed.

**RESULTS:** Compared to N-S, LPS-stimulated monocytes display a singular milieu of markers, with higher levels of intracellular IL-1β in parallel raise of CD14+CD163-/CD14+CD163+ ratio. CD163 shows positive correlation with levels of IL-1β. In t-SNE (T-distributed Stochastic Neighbor Embedding) analysis, after LPS stimulation, subpopulation CD14+CD16-CD163-, containing mainly classical monocytes, show a higher number of IL-1β+ cells.

**CONCLUSION:** Classical monocytes, the non-inflammatory subset, show higher levels of IL-1β in response to LPS narrowing down to a new subpopulation of monocytes CD14+CD16-CD163-, which correlates better with this interleukin response than widely used monocytes classification. Using CD163 in addition to CD16, we were able to show that classical monocytes display the most intense response to LPS. Additionally, CD163 appears to be a suitable addition to CD14-CD16 classification to improve its performance.

## Introduction

In 2010, an international meeting sponsored by the IUIS (International Union of Immunological Societies) and the WHO (World Health Organization), proposed a subset classification for circulating monocytes (1). CD14 (Monocyte identifying Toll-Like Receptor) and CD16 (FcyRIII co-receptor, marker of inflammatory monocyte) were used to define 3 subpopulations of circulating monocytes with different attributes in terms of inflammatory protein expression and phagocytic capabilities (2). Classical monocytes (CD14++CD16−) are 80 to 90% percent of the entire monocyte’s population, express low levels of pro-inflammatory proteins and high phagocytic characteristics. Intermediate monocytes (CD14+CD16+) are 5 to 15% of the total monocytes. Express variable levels of pro-inflammatory cytokines and intermediate phagocytic capacities. Non-classical monocytes (CD14+CD16++) (3) are approximately 5% of all monocytes, express higher levels of pro-inflammatory cytokines and minor phagocytic characteristics. The last two subsets, since have shown to express higher levels of inflammatory cytokines when stimulated with pro inflammatory stimuli, have been repeatedly related to inflammatory pathologies. (4).

During the last years, multiple studies have been published associating expansion or contraction of inflammatory or non-inflammatory monocytes subpopulations in diseases like lupus, rheumatoid arthritis, atherosclerosis or membranous glomerulonephritis (5–7), among others. Intermediate monocytes has shown to be expanded in infectious and non-infectious inflammatory conditions, however many studies presents inconsistent and contradictory data.(8).

In cardiovascular diseases, intermediate monocytes have been reported both expanded and contracted(8, 9). In the other hand, Classical monocytes show higher frequencies in atherosclerotic lesions despite of inflammatory status, evidencing conflicting data in this regard(9, 10). In 2017, Williams et al. classified circulating monocytes from post-stroke patients, using CD86 (co-stimulatory T signal, typically described M1 Monocyte/Macrophage phenotype) and CD163 (High-affinity scavenger receptor typically related with M2 Monocyte/Macrophage phenotype) to group monocytes as M1 and M2 trying to emulate macrophage classification. They showed that M1/M2 ratio (CD86/CD163 ratio) was higher in post-stroke patients(11, 12). In other clinical entities like membranous glomerulopathy, similar results have been found using broader group of markers defining M1 and M2 phenotypes (5).

Until now, many changes have been introduced looking for accuracy to define monocytes subpopulations. Thomas et al. using multiparametric analysis, tSNE (t-distributed stochastic neighbor embedding), showed that CCR2 (CC Chemokine Receptor Type 2), HLA-DR (Human Leukocyte Antigen – DR isotype) and CD163 were good and enough markers to a proper M1 and M2 monocytes classification (10). Interestingly CD163 and CD16, protein which defines current monocyte classification, does not show correlation in clinical setting (13).

In vitro, when monocytes are stimulated with pro-inflammatory and anti-inflammatory stimuli, there are also contradictory reports in terms of interleukin production response. Mukerjhee et al. showed that monocytes from healthy donors do not display cell number changes when stimulated with LPS. Also, intermediate and non-classical monocytes express higher levels of IL-1β and TNF-α, but inflammatory intermediate subpopulation, responds with higher levels of regulatory IL-10 to the same stimulus (7). Subsequent studies have shown that M1 monocytes express higher levels of IL-6 when stimulated with LPS exposing again contradictory reports(11).

Given the absence of a solid classification strategy and in vitro monocyte response to inflammatory stimuli, we analyzed monocytes from nine healthy donors and expose it to LPS and IL-4 showing a singular cytokine production and phenotypic pattern. We found that less-reactive-non-inflammatory classical monocyte subpopulation has the most robust response *in vitro* to LPS stimulus. This subpopulation also could be better classified using other membrane markers as CD163. Classical monocytes show also a more “flexible” M1/M2 ratio, compared to intermediate and non-classical monocytes, becoming the subpopulation with the highest functional plasticity.

## Material and Methods

### 2.1. Healthy donors

We recruited 9 healthy donors ranging from 26 to 40 years, 5 man and 4 women, from our facility. A medical anamnesis was performed by an internal medicine specialist before whole blood extraction. No medication uses nor relevant past medical history were reported. The ethics committee of Facultad de Medicina de la Universidad de Chile approved the study protocol.

### 2.2. Whole blood lysis protocol and flow cytometry

Venous blood (30ml) was obtained by cubital venopunction from all participants. PBMC Ficoll extraction was performed as previously dercribed(REF). PBMC were stained in duplicate with the following antibodies: FITC-A-anti-CD14, APC-H7-anti CD16, PERCP-Cy5.5-anti-CD86, APC-A-anti-CD16, V450-A-anti-IL-1β (Biolegend, US) at room temperature for 30 minutes. 1ml of BD FACS Lysing Solution (BD Biosciences) at 1x concentration was used for erythrocyte lysis. Finally cells we fixed and permeabilized using a fixation/permeabilization BD Kit (BD Biosciences).

### 2.3. LPS and IL-4 Stimulation and flow cytometry

To phenotype monocyte subsets, 1 × 10^6^/cells per tube were stimulated in duplicate with lipopolysaccharide, (1 µg/mL) (BD Biosciences, NY) and IL-4 (200 U/ml) (BD Biosciences, NY) in RPMI 1640 (BD Biosciences, NY) medium mixed with 10% fetal bovine serum (BD Biosciences, NY) for 2 h and 4 h respectively at 37°C in 5% CO2 and exposed to brefeldin A (BD Biosciences) as described in previous studies. Then, the cells were washed, followed by staining with V710-anti-CD3 (Biolegend, US), V650-anti-CD19, (Biolegend, US), FITC-A-anti-CD14, APC-H7-anti CD16, PERCP-Cy5.5-anti-CD86, APC-A-anti-CD16, (Biolegend, US). Subsequently, they were fixed and permeabilized and stained with V450-A-anti-IL-1β (Biolegend, US). The frequencies monocyte subsets were assessed on a FACS Verse instrument (BD, Franklin Lakes, US), and the data was analyzed with FlowJo software (v7.6.1; TreeStar; Ashland, US).

### 2.4. Statistical Analysis

Variables were shown as median and range values. We employed the Mann–Whitney U nonparametric test to evaluate the differences among groups. The relationship between variables was analyzed by the Spearman rank correlation test. All of the data were carried out with the GraphPad Prism version 6.01 software. P value of <0.05 represented statistical significance.

## Results

### Classical monocytes are the most responsive subpopulation to LPS stimuli measured as IL-1β MFI

Monocytes where gated using CD14 and CD16 for usual three-subsets classification (Fig1, A, B, C). Classical monocytes are significantly higher than intermediate and non-classical monocytes in any condition and percentages are equivalent to previously described studies(1) (Fig1, D). In each subpopulation, levels of IL-1β measured as Mean Fluorescence Intensity (MFI) and count of IL-1β+ monocytes, are higher when stimulated with LPS, being significantly higher in Classical versus Non-classical monocytes (Fig1 E, F).

**FIGURE 1:**
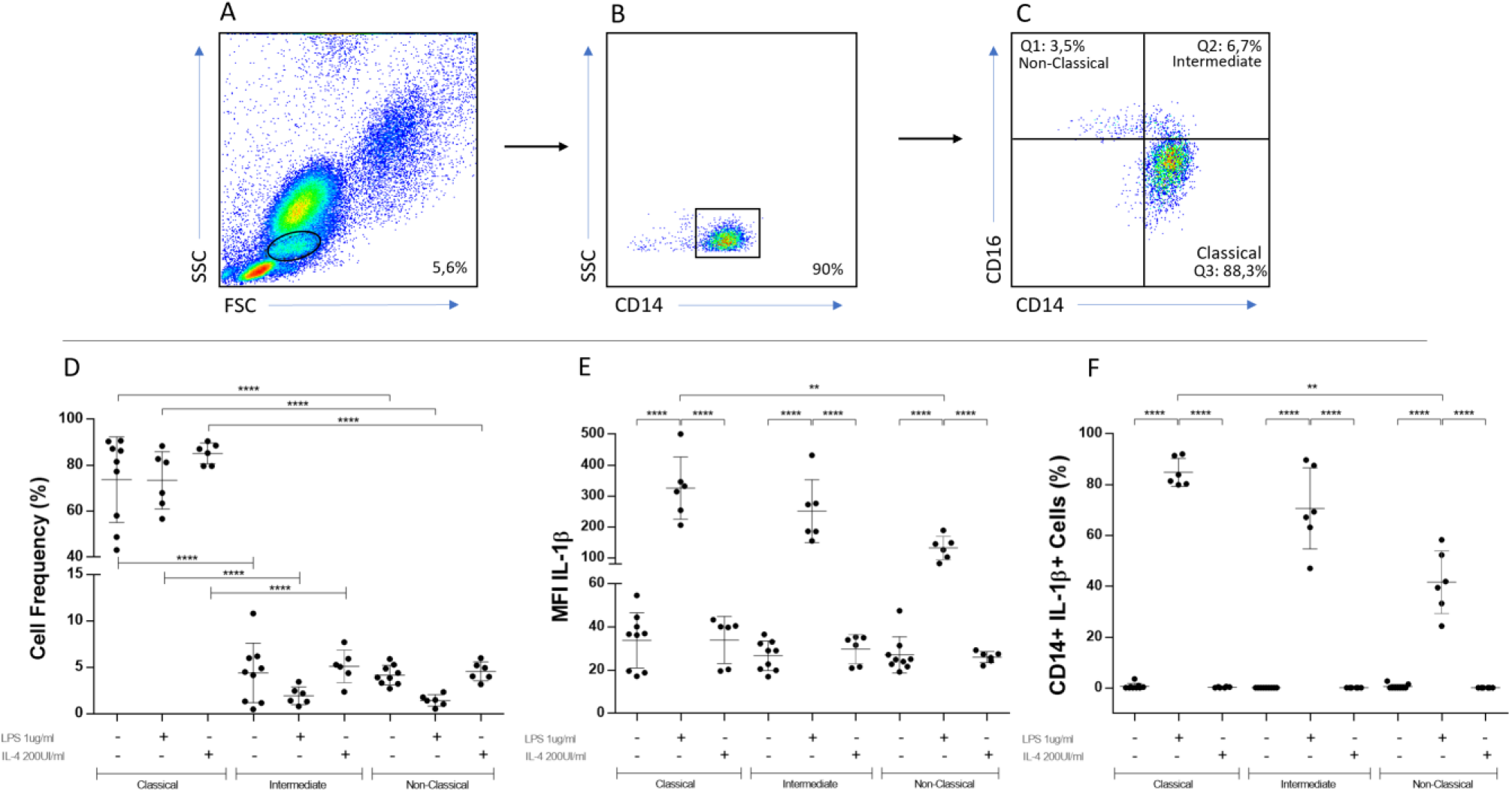
Gating strategy, flow cytometry analysis and intracellular IL-1β in CD14-CD16 defined subpopulations under different stimulation conditions. Whole blood from healthy donors was stained with anti-CD14, anti-CD16, anti-CD86 and anti-CD163. The cells were gated firstly on mononuclear cells (A) and secondly on CD14+ monocytes (B). Afterwards, the count of CD14++CD16-, CD14+CD16+ and CD14+CD16++ were determined by flow cytometry (C). Classical monocytes are significantly higher than intermediate and non-classical monocytes in any condition (D)(****p≤0,0001). In each subpopulation, levels of IL-1β are higher when stimulated with LPS, being significantly higher in Classical versus Non-classical monocytes (E) (****p≤0,0001; **p≤0,007). When IL-1β+ cells are analyzed, LPS stimulated monocytes, show significantly higher cell count in Classical versus Non-Classical monocytes (F) (**p≤0,009).

### CD163- monocytes rise in response to LPS and shows higher levels of intracellular IL-1β

Monocytes were firstly gated using CD14 and then, using CD163, two different subpopulations where found with evenly split frequencies (Fig2 A, B, C).When expose with LPS, CD163- cells count raises to a maximum reciprocally to CD163+ cells which shows lowest levels. IL-4 monocytes maintain 50%-50% proportion seen on basal conditions (Fig2, D). CD163- cells contains significantly higher levels of IL-1β than CD163+ monocytes (Fig2, E). When CD14+CD163-/CD14+CD163+ Ratio is analyzed, the abrupt rise in CD163- cells seen when expose to LPS, reflects as a higher ratio only in Classical monocytes, suggesting higher plasticity on this subpopulation (Fig2, F).

**FIGURE 2:**
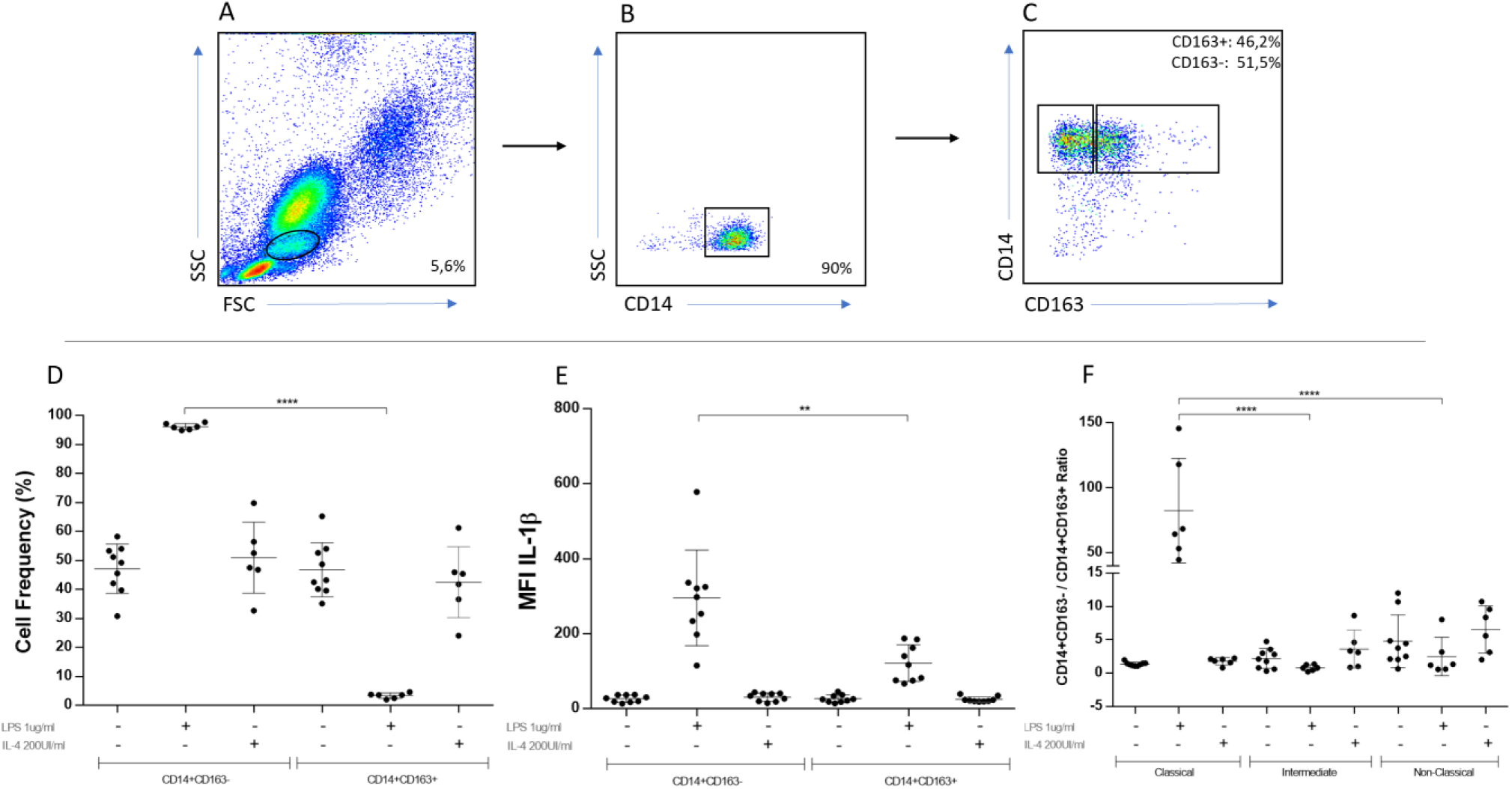
Gating strategy, flow cytometry analysis, intracellular IL-1β in CD14-CD163 defined subpopulations under different stimulation conditions. Whole blood stained with anti-CD14, anti-CD16, anti-CD86 and anti-CD163. The cells were gated firstly on mononuclear cells (A) and secondly on CD14+ monocytes (B). Afterwards, the count of CD14+CD163- and CD14+CD163+ were determined by flow cytometry (C). CD163- cells count raises only when stimulated with LPS, reciprocally to CD163+ cells (D)(****p≤0,0001). CD163- cells contains significantly higher levels of IL-1β than CD163+ monocytes (E)(**p≤0,009). Only in Classical monocytes stimulated with LPS, CD163-/CD163+ ratio raises (F)(****p≤0,0001).

### Only CD163 and no CD16, show correlation with IL-1β after LPS stimulation

When we analyzed separately positivity to either CD163 or CD16 and levels of IL-1β after LPS stimulation, only CD163 negativity pairs with IL-1β peaks in all subsets (Fig3, A). After correlation tests, again only CD163 show positive and significant correlation with MFI IL-1β after LPS stimulation (Fig3, B, C). No correlation is seen between CD16 and IL-1β (Fig3, D, E).

**FIGURE 3:**
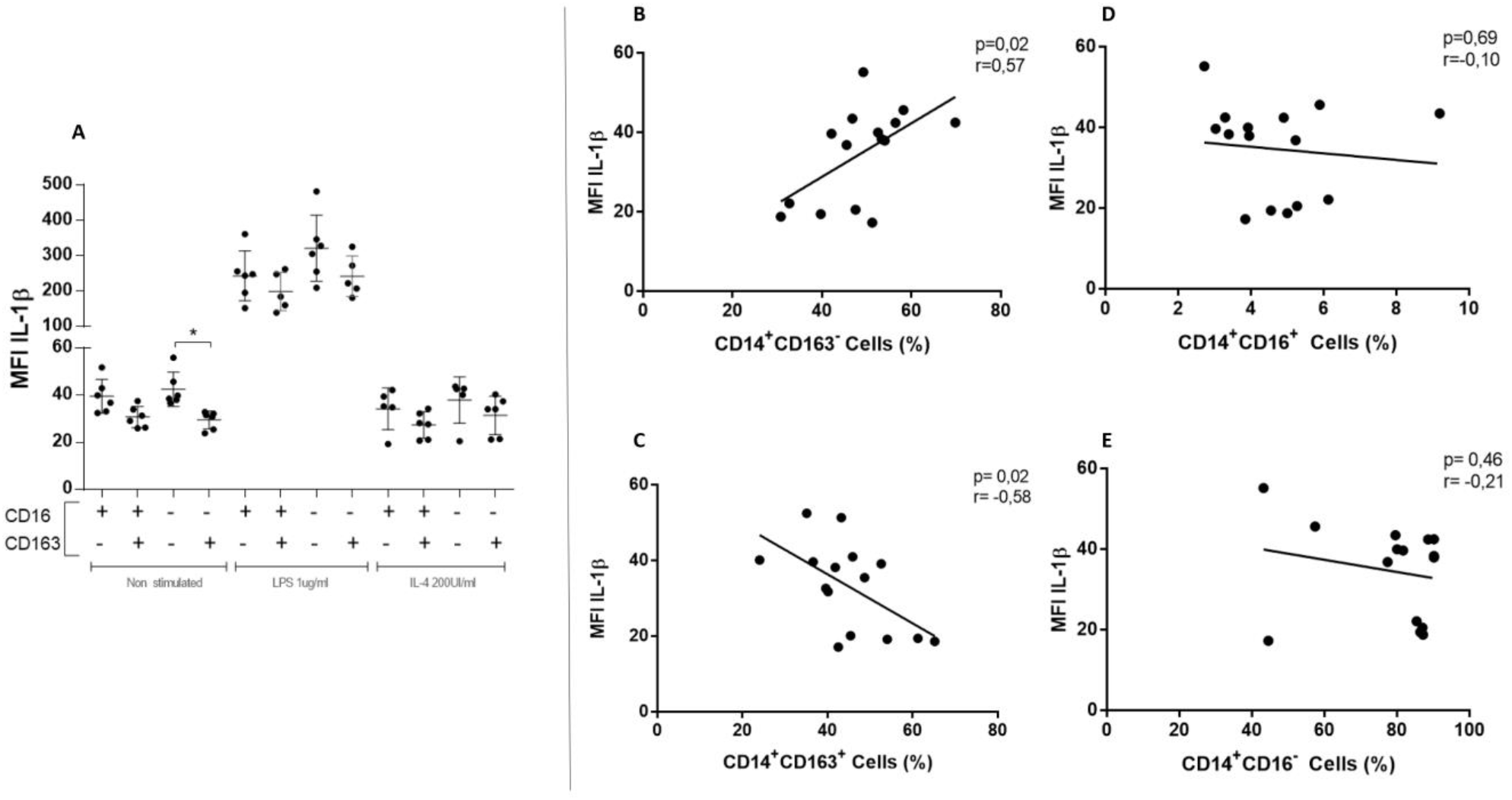
Intracellular IL-1β correlation with both CD16 and CD163 markers. Association between levels of IL-1β and CD16-CD163 positivity/negativity in each stimulation condition. In A, positivity for CD16 marker, did not show relation with levels of IL-1β. On the contrary, negativity for CD163 is present every time IL-1β raises, being significantly higher in N-S condition (*p≤0,02). Correlation between CD163 and CD16 cells and levels of intracellular IL-1β. CD14+CD163- cells correlate positive and significantly with levels of IL-1β, on the contrary CD163+ have negative correlation (B, C). CD16+ o CD16- cells did not correlate significantly with levels of intracellular IL-1β (D, E).

### CD14+CD163- inflammatory subset rather CD14+CD16+ subset, respond to LPS stimulation raising intracellular IL-1β

Using t-SNE multiparametric reduction (of CD3, CD19, CD14, CD16, CD163, CD86 markers), we were able to reduce all monocytes to two cell clouds, CD14+CD16+ and CD14+ CD163- which interestingly, where mutually exclusive (Fig4, A). When stimulated with LPS, IL-1β+ monocytes invade CD163- cells without significantly grow on CD16+ cloud, showing that CD163- monocytes are especially responsive shaping a new inflammatory subpopulation CD14+CD16-CD163-IL-1β+ (Fig4, C).

**FIGURE 4:**
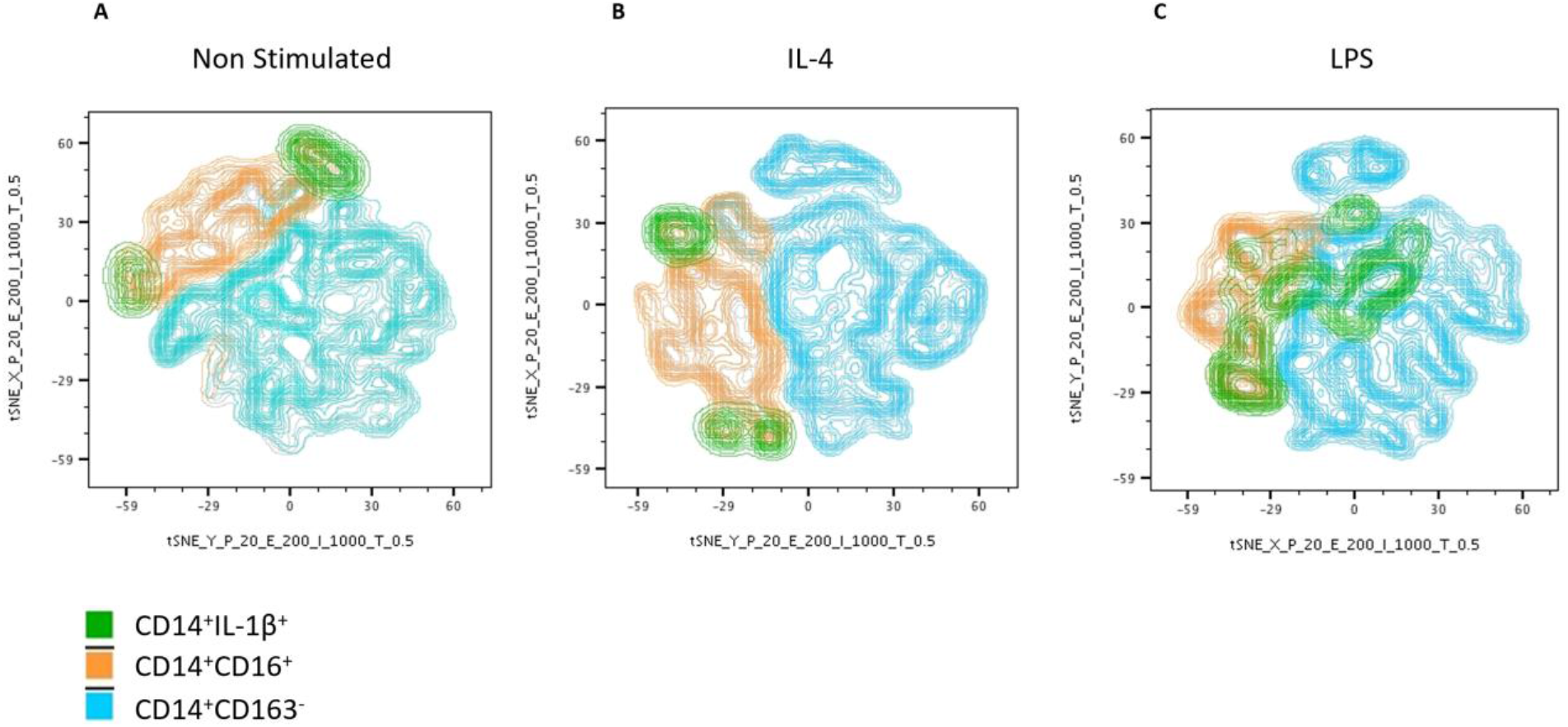
t-SNE analysis for CD14+ monocytes using multiple markers. Orange colored cloud of cells represents CD16+ cells, light blue cloud CD163- cells and green cloud IL-1β+ cells, being the first two, “inflammatory” subpopulation according most used classifications. As is easily seen, CD16+ and CD163- cells are completely different subpopulations (A). In IL-4 stimulation condition, no changes are seen in green cloud, which is comparable to N-S condition (B). In C, after LPS stimulation, green cloud “invades” CD16-CD163- forming a new subpopulation CD14+CD16- CD163-IL-1β+ (C).

## Discussion

During the last years, numerous studies have tried to find an association between certain monocyte subpopulations and prevalent inflammatory and metabolic diseases. Autoimmune entities and atherosclerosis as a metabolic dysregulation are the most usually studied, nevertheless, their correlation with expansion or contraction of these subpopulations has been difficult to study and not completely reproducible. Many reasons for the latter have been proposed. One of the most frequently argued explanation is how difficult is to separate three “different” subpopulations using just a small group of markers. Many times, depending on the inflammatory status of the patient, the first selection criteria in the gating process, as is the FSC (granularity) in flow cytometry, could represent a challenge trying to gate only monocytes out of a “universe” of neutrophils.

Thomas et al. were able to demonstrate using tSNE with more than 20 different parameters, that a group of at least 12 markers commonly available, must be used to perform positive and negative selection in order to obtain a proper separation (14). Even when these markers help to better classify circulating monocytes, it is yet expensive and time consuming to process it.

When these monocyte subpopulations are analyzed *in vitro*, its response to stimuli turns more erratic. Most of the studies available, are prone to point to intermediate monocyte subpopulation as the most “plastic” and responsive to LPS. Many times, intermediate monocytes are presented as transition subpopulation with oscillations on percentages attributed to inflammatory nature on each pathology. Despite all this, until today, there is no clear correlation between any subpopulation rate changes and clinical course in inflammatory or non-inflammatory conditions.

In our experience, we were able to show that non-inflammatory classical monocytes constitute the most responsive subpopulation. Classical monocytes uniquely raise levels of intracellular IL-1β. Also, when these monocytes are reclassified using CD163, CD163- cells are the most responsive. When compared CD16 marker and CD163 marker, the last is the one who better correlate with intracellular levels of IL-1β. Classical monocytes subset, a priori classified as less-responsive/non-inflammatory subset, contains almost 100% of high responsive CD163- cells.

We were also able to show that, in the way of a Russian *matryoshka*, classical monocytes display an intern and variable plasticity, with different grades of responsiveness depending at least in part, on CD163- cells number. These findings rise the question about if we are properly classifying monocytes subsets using a bidimensional analysis. More studies are needed to understand if the most responsive subpopulation CD14+CD163- and a new subpopulation CD14+CD16-CD163-IL-1β+ displays different sub-phenotypes with also different grades of response to LPS and other stimuli using more markers or multiparametric analysis. How these sub-phenotypes behave in clinical scenarios remain a question to be investigated.

